# Analysis of Finnish blue mussel (*Mytilus edulis* L.) shell: Biomineral ultrastructure, organic-rich interfacial matrix and mechanical behavior

**DOI:** 10.1101/636696

**Authors:** Pezhman Mohammadi, Wolfgang Wagermaier, Merja Penttila, Markus B. Linder

**Affiliations:** VTT Technical Research Centre of Finland Ltd., Espoo, FI-02044, VTT, Finland; Department of Biomaterials, Max Planck Institute of Colloids and Interfaces, D 14424 Potsdam (Germany); Department of Bioproducts and Biosystems, School of Chemical Engineering, Aalto University, P.O. Box 16100, FI-16100, Espoo (Finland)

**Keywords:** Finnish blue mussel, Biomineralization, Interfacial matrix protein, Chitin

## Abstract

Studying various marine biomineralized ultrastructures reveals the appearance of common architectural designs and building blocks in materials with fascinating mechanical properties that match perfectly to their biological tasks. Advanced mechanical properties of biological materials are attributed to evolutionary optimized molecular architectures and structural hierarchy. One example which has not yet been structurally investigated in great detail is the shell of *Mytilus edulis* L. (Finnish blue mussel) found in the archipelago of SW-Finland. Through a combination of various state-of-the-art techniques such as high-resolution electron microscopy imaging, Fourier-transformed infrared spectroscopy, powder X-ray diffraction, synchrotron wide-angle X-ray diffraction, nanoindentation and protein analysis, both the inorganic mineralized components as well as the organic-rich matrix were extensively characterized. We found very similar ultra-architecture across the shell of *M. edulis* L. as compared to the widely studied and closely related *M. edulis*. However, we also found interesting differences, for instance in the thickness and degree of orientation of the mineralized layers indicating dissimilar properties and related alterations in the biomineralization processes. Our results show that the shell of *M. edulis* L. has a gradient of mechanical properties, with the increase in the stiffness and the hardness from anterior to the posterior region of the shell. The shell is made from distinct and recognizable mineralized layers each varying in thickness and microstructural features. At posterior regions of the shell, moving from dorsal to ventral side, these layers are an oblique prismatic layer, a prismatic layer and a nacreous layer, in which the oblique prismatic layer is found to be the main and thickest mineralized layer of the shell. Probing the calcified rods in the oblique prismatic layer using high resolution SEM imaging revealed opening of channels with a diameters of 40-50 nm and lengths up to a micrometer extending through the rods. The chitin and protein have been found to be the main component of the organic-rich interfacial matrix as expected. Protein analysis showed two abundant proteins with sizes around 100 kD and 45 kD which likely not only regulates biomineralization and adhesion of the crystals but also governing the intrinsic-extrinsic toughening in the shell. Overall, this detailed analysis provides new structural insights into biomineralization of marine shells in general.

## 1. Introduction

Biological materials show fascinating properties and in many ways can outperform man-made engineered materials, although they are built from very limited elements appearing in nature and comparable simple building blocks ^1^. Exceptional examples of such materials can be found in marine biomineral structures that combine high stiffness, strength, and toughness. Such materials achieve their function by assembling large fractions of anisotropic inorganic stiff and strong elements, embedded in an isotropic soft and energy-dissipating adhesive organic matrix ^2^. These two basic components with mismatching properties are merged in a complex hierarchical architecture, which ultimately enables absorption of energy at different length scales. This is driven by evolutionary optimized molecular interactions among various building blocks throughout the course of the evolution.

For instance, nacre in the interior of mollusk shells -also known as mother of pearl-features a brick and mortar microstructure, assembled from 0.4-0.5 μm thick and 5-15 μm wide tablets of calcium carbonate (aragonite) that are glued to one another with few nanometer thin continuous layers of adhesive and energy dissipating organic matrix ^3,4^. Orders of magnitude tougher than nacre, conch shell is another example of a marine biomineral structure with a three-tier multiscale lamellar structure ^5–7^. Shells consist of outer, middle and inner layers, each further constitute of first (5 μm thick and several μm wide), second (5-30 μm thick and 5-60 μm wide) and third (60-100 nm thick and 100-400 nm wide) order lamellae varying in thickness and length of the aragonite platelets that are glued to one another by the organic matrix. This hierarchical, lamellar ultra-architecture provides various crack deflecting pathways, bridging and fiber pullout, hence increasing the toughness.

Details of assembly mechanisms and the formation of such complex hierarchical structures are still under investigation. However there is growing evidence illustrating that the organic-rich interface consisting of silk-like and acidic proteins along with a chitin network ^8,9^, largely governs both nucleation and inhibition of the crystal growth and providing a matrix structure that has substantial importance toward structural integrity of the shell ^10,11^.

In recent years, biological materials have been studied to understand basic principles of biomineralization making those available for material synthesis as well as one main source of inspiration for fabrication of next generation high performance and advanced materials (Studart, 2016, 2012; Wegst et al., 2015; Xu et al., 2007). These potential applications range from advanced medical devices, robust electronics apparatus, and sensors, 15–17 to lightweight aerospace application and military armors ^18,19^.

Despite its scientific, ecological and economic importance, the current understanding of such complex mineral producing machinery is far from complete. This motivates to analyze the ultrastructure and mechanical properties of biomineralized materials which have never been studied before to identify common or dissimilar architectural features and shade light onto outstanding mechanical characteristics of such materials. Furthermore, a fundamental understanding of structure-property relations has substantial implications for design strategies and manufacturing processes of next-generation advanced functional materials ^20^.

In this study, we provide extensive characterization of the shell of *M. edulis* L. (Finnish blue mussel) found in the archipelago of SW-Finland (Northern Baltic Sea) for the first time. This was carried out by combing various state of the art techniques to study both inorganic mineralized components as well as the organic matrix, including high-resolution electron microscopy imaging, Fourier-transform infrared spectroscopy (FTIR), RAMAN-spectroscopy, powder X-ray diffraction, synchrotron wide-angle X-ray diffraction (WAXS), Nanoindentation and proteomic analysis.

## 2. Method and materials

### 2.1. Research specimens

Live specimens of *M. edulis* L. were collected from the archipelago of SW-Finland in the northern Baltic Sea. Shells were dissected and removed. The outer sides of the shells were cleaned mechanically for removing contaminants and epibionts. The inner sides of the shells were also cleaned from connective tissue and rinsed with cold (4 °C) water to remove any loose organic debris. Samples were immediately flash-frozen in liquid nitrogen and kept at −80°C in a freezer until use.

### 2.2. Scanning electron microscopy (SEM)

SEM imaging carried out with a Zeiss FE-SEM field emission microscope (Microscopy center, Aalto University, Espoo, Finland) with variable pressure, operating at 1–1.5kV operating in low vacuum mode for imaging hydrated specimens.

### 2.3. Powder X-ray diffraction (XRD)

XRD carried out using Rigaku SmartLab, equipped with HyPix-3000 Hybrid Pixel Array Detector. Collection performed using CuKα-radiation of λ=1.5418 Å (energy of 45 kV and 40 mA). The diffractometer collected at a 2θ range of 10-80° with the step size of 0.01°/s and 4 sec exposure time. The crystalline phases were identified by matching the XRD patterns with the Joint Committee on Powder Diffraction Standards (JCPDS) database.

### 2.4. Synchrotron wide-angle x-ray scattering (WAXS) measurement

Wide-angle X-ray diffraction experiments were performed at the μSpot beamline at BESSY II synchrotron source (Helmholtz-Zentrum Berlin für Materialien und Energie, Germany). The measurements were carried out using a silicon (111) monochromator with an X-ray wavelength of 0.82656Å (energy of 15 keV) and a beam size of 50 μm. The beamline calibration was done with a quartz (SiO_4_) standard giving a sample to detector distance of approximately 280mm. Diffraction patterns were collected using a two-dimensional CCD detector (MarMosaic 225, Mar USA, Evanston USA) with a pixel size of 73.242 μm and 3072 × 3072 pixels. Intensities have been characterized using the software DPDAK after subtraction of air scattering from the diffractogram ^21^.

### 2.5. Nanoindentation

Nanoindentation was performed under ambient conditions using an Ubi nanoindentation instrument (Hysitron) with a Berkovich diamond tip. To obtain reduced modulus and hardness, the load-displacement curves were analyzed using the methods described by Oliver and Pharr. ^22^ More than 1000 indentations were performed per sample using a maximum load of 1500 μN on the polished surface. At peak load, a dwell time of 10 s was applied to account for creep behavior. At five different regions across each sample, two adjoined lines were measured with steps of 15 μm (between lines) and 6.2 μm between each indent along the line. Therefore, plastically deformed zones from previous indents did not affect the measurements. The effect of the topography was minimized by finely polishing the sample surface.

### 2.6. Fourier transform infrared spectroscopy (FTIR)

Infrared measurement carried out using Unicam Mattson 3000 FTIR spectrometer directly on the finely powdered shell. To do that, the specimen was mixed with potassium bromide (1:9 mixing ratio of the specimen in KBr) which was then turned into 1mm transparent discs under high pressure and measured in transmission mode. All spectra were scanned within the range of 400–4000 cm^-1^, with a total of 32 scans and a resolution of 32 cm^-1^.

### 2.7. Protein extraction

Shells were incubated with 1%, v/v NaOCl for 24 hours at 4 °C in order to remove superficial organic contaminants as well as the periostracum. Shells were then rinsed with deionized water and dried at ambient temperature before mechanically crushing them into powder in a cold room (4 °C). Decalcification carried out by addition of 50 ml 1 M EDTA pH 8 to every 5 g of shell powder for 24 hours at 4 °C and centrifuged at 40000g for 60 min. Supernatants collected and dialyzed against MQ water and then freeze-dried. Pellets from each method also rinsed three times with MQ water and freeze-dried. Extracted proteins separated and analyzed with standard sodium dodecyl sulfate-polyacrylamide gel electrophoresis (SDS*-*PAGE) ^23^.

### 2.8. MALDI-TOF-TOF

Proteins identity was confirmed by matrix-assisted laser desorption/ionization-time of flight/time of flight (MALDI-TOF-TOF) mass spectrometer (UltrafleXtream™ Bruker, Aalto department of biotechnology and chemical technology facilities, Espoo, Finland) equipped with a 200-Hz smart-beam laser (337 nm, 4 ns pulse). For partial peptide sequencing (MS/MS), in-gel tryptic digestion performed using proteoprofile trypsin in-gel digest kit (PP0100, Sigma-Aldrich). The samples were desalted using either ZipTip™ C18 (Millipore) eluted directly onto MALDI target plate using MALDI matrix (a-cyano-4-hydroxycinnamic acid, 10 mg/ml in 70% ACN, 0.03% TFA). Spectra acquired in the reflection positive ion mode and resulting monoisotopic masses cross-referenced against the database (NCBI) with a tolerance of 150 ppm. Protein identification performed using the MASCOT search engine (Matrix Science, London, UK; version 2.1) against NCBI server (http://www.ncbi.nlm.nih.gov). The peptide mass and fragment ion tolerances were set to 0.5 Da for MS/MS data searched. The peptide peaks further manually investigated by the interpretation of the raw MS/MS spectra to perform *de novo* sequences. Furthermore resulted peptides cross-referenced against BLASTp, Swiss-Prot, and UniProt databases.

## 3. Results and discussion

### 3.1. Research specimen

Selected *M. edulis* L. obtained from the archipelago of SW-Finland in the northern Baltic Sea for this study had a size of about 2.5-3 cm (anterior-posterior length) (Fig. 1A, B, and S1). In general, the size of the adult *M. edulis* L. is relatively smaller than other extensively studied and closely related taxa of mussels such *M. edulis* (commonly found in North Atlantic and Pacific), *M. galloprovincialis* (Mediterranean, black sea, and North Pacific) *and M. trossulus* (North Atlantic and Baltic Sea). Mytilus specimens in our study have been found to be a crossbreed between *M. edulis and M. trossulus* ^24^.

**Fig. 1.**
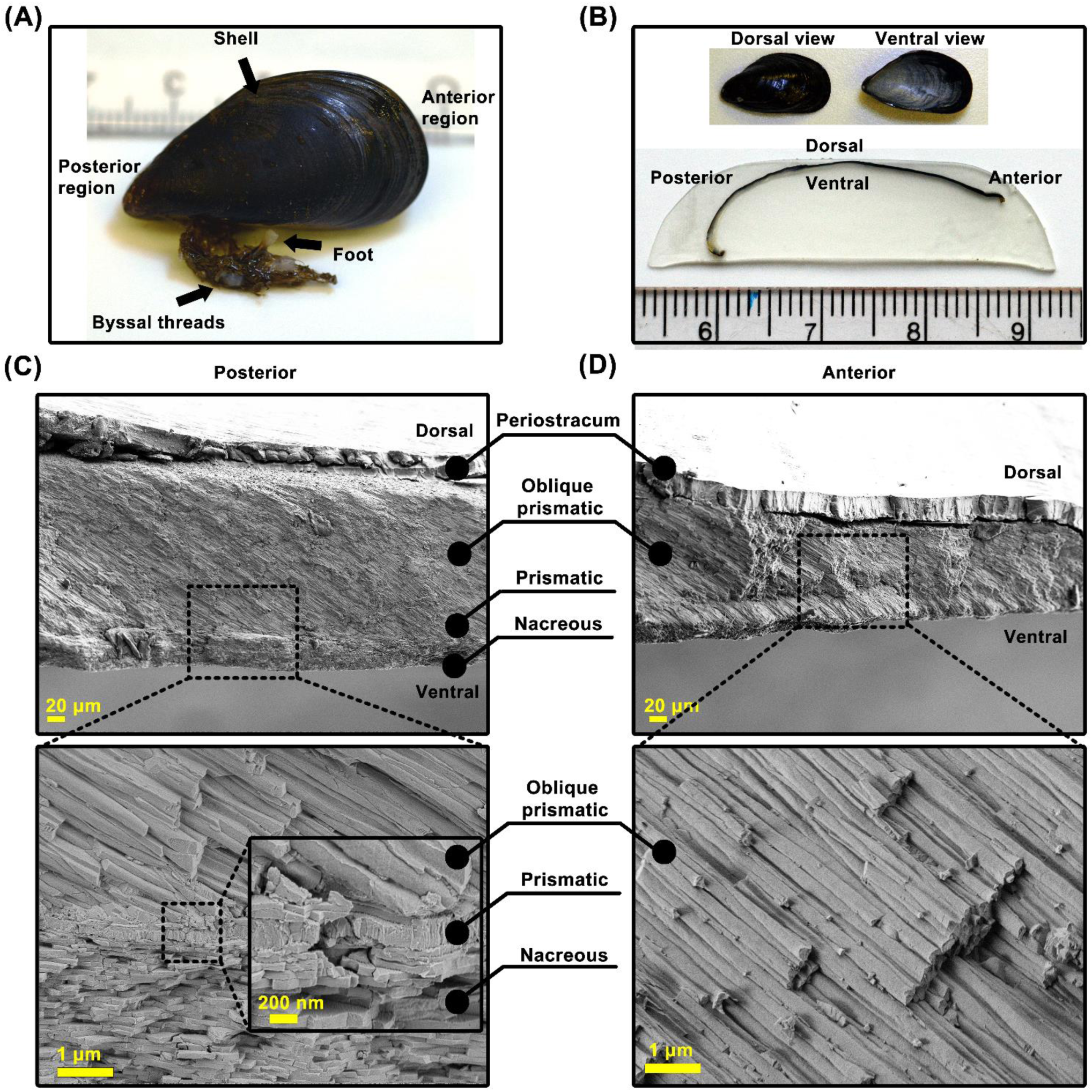
(A) Morphological features of an adult M. edulis L. shell, commonly known as Finish blue mussel obtained from the archipelago of SW-Finland in the northern Baltic Sea. (B) Longitudinal cross-section of the shell. (C and D) SEM micrographs from the cross-section of the shell taken from posterior and anterior regions at different magnifications. SEM images illustrating periostracum layer, oblique prismatic layer, prismatic layer and a nacreous layer which can be found at the posterior region. At the anterior region, only periostracum and oblique prismatic layers can be found.

### 3.2. Biomineral ultra-architecture of the shell

Scanning electron microscopy (SEM) of cross-sections of the shell revealed presence of multiple layers, however distinct and recognizable mineralized layers each varying in thickness and microstructural features. At the posterior regions of the shell, from dorsal to ventral side, the outer most layer starts with an organic layer called periostracum. Periostracum is a protective leathery layer that has a substantial importance for the mineralization and templating of the first calcified layer. Subjacent to the periostracum is the calcified oblique prismatic layer, exhibiting a closely packed, ordered and highly elongated rod-like ultrastructure, which is developed around 19° relative to the periostracum layer. Subsequently, the prismatic and nacreous layers are located, both developed perpendicular to the periostracum (Fig. 1C). The prismatic layer consists of radially elongated short monocrystals while the interior nacreous layer comprises a brick and mortar microstructure, assembled from tablets with uniform thickness and widths. Among all, oblique prismatic sheets represent the main and thickest (≃ 200 μm) mineralized layer. The oblique prismatic layer alone constitutes about 86% of the overall shell thickness in comparison to the nacreous (≃ 30 μm) and prismatic (≃ 0.4 μm) layer which only represent about 13% and less than 0.05 % respectively. In addition, probing the cross-section of the shell at the anterior region (growth front) revealed the presence of only the periostracum and oblique prismatic layer (with a thickness of ≃125 μm forming 100% calcified layer) (Fig. 1D). We found many similarities in the ultra-architecture of the *M. edulis* L. shell in this study and the closely related *M. edulis*. However, we also found differences in the thickness of the layers comparing these result to an earlier studied adult *M. edulis* shells ^25^. Table 1 summarizes these differences.

**Table 1.** M. *edulis* L. versus *M. edulis* shell. Comparison between the thickness of inorganic and organic layers in the M. edulis L. and M. edulis shell.

| Table 1. *M. edulis* L. versus *M. edulis* shell |  |  |  |  |
| --- | --- | --- | --- | --- |
| Layers | *M. edulis* L. |  | *Mytilus edulis* |  |
|  | μm | % | μm | % |
| **Inorganic** |  |  |  |  |
| Oblique prismatic | 200 | 86 | 296 | 80 |
| Prismatic | 0.2 | 0.05 | 4 | 1 |
| Nacreous | 30 | 13 | 66 | 18 |
| **Organic** |  |  |  |  |
| Periostracum | 30 | - | 15 | - |

### 3.3. Calcite and aragonite

To investigate the nature of the shell and its variability in the calcified composition we performed XRD. The results (Fig. 2A) illustrated that the shell, in fact, contains calcium carbonate (CaCO_3_) which has been shown for many other seashells ^26^. Spectra showed that the shell mainly constitutes of calcite, but reflections corresponding to aragonite were also noticeable. Calcite and aragonite both were found to be the most stable polymorph of CaCO_3_. We further mapped the distribution of each polymorph across the shell using simultaneous high-resolution synchrotron WAXS/SAXS mapping (Fig. 2B and C). This revealed two very distinct phases of calcite and aragonite. A thick outer region comprises calcite corresponding to the oblique prismatic layer and a thinner region toward the ventral side comprise of aragonite corresponding to prismatic and nacreous layers.

**Fig. 2.**
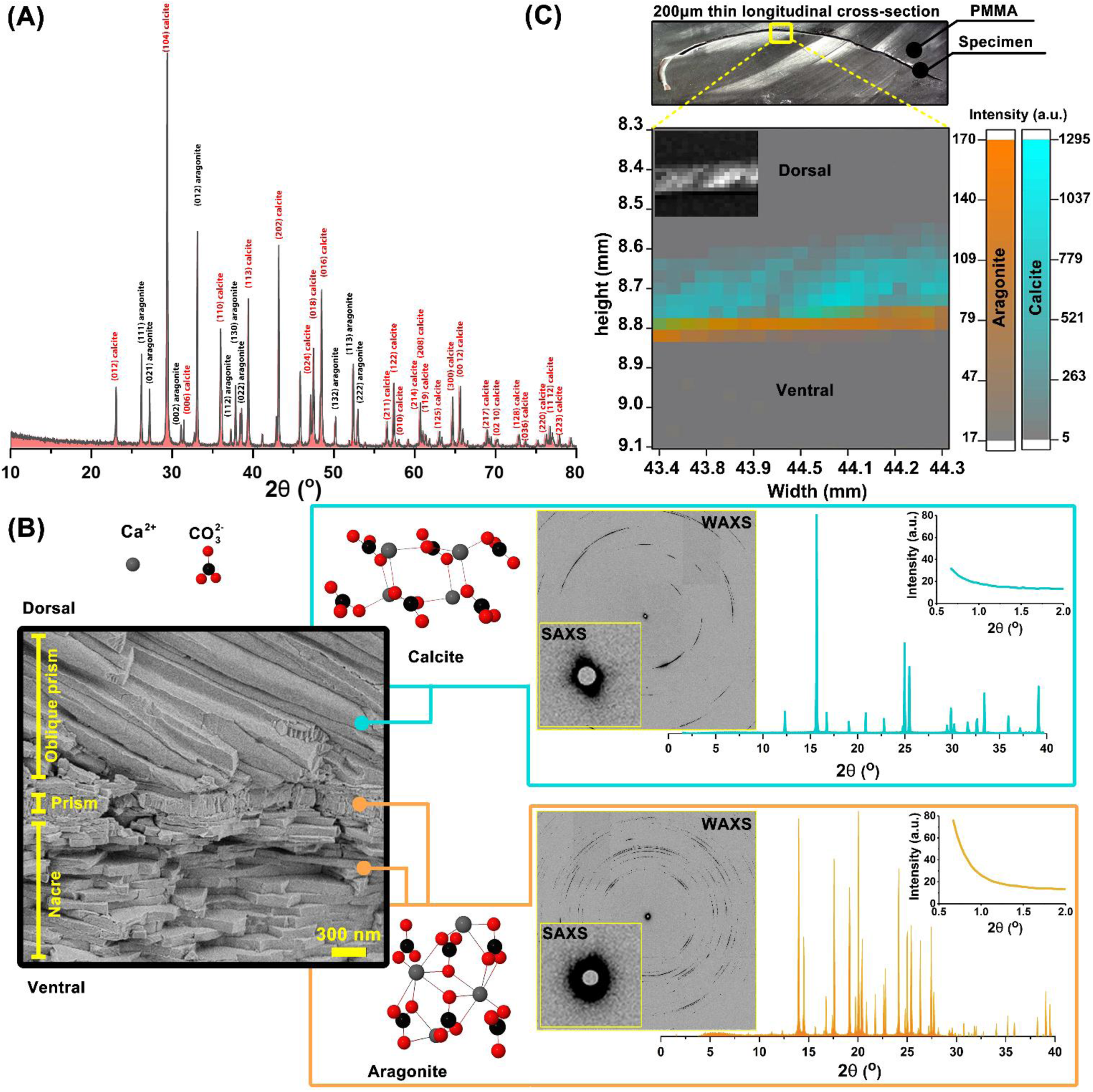
(A) XRD of the bulk shell powder. (B) Representative 2D synchrotron WAXS/SAXS diffraction patterns and their extracted 1D profile corresponding to oblique prismatic and nacreous mineralized layers. (C) Heat-map illustrating scattering intensity distribution of the calcite and aragonite throughout the ultra-architecture of the shell. Each square corresponds to single measurement point. Cyan indicates distribution of the calcite and orange displays the aragonite.

### 3.4. Oblique prismatic layer

High magnification SEM images from oblique prismatic showed that it is made from individual calcified rod-like ultrastructure with diameter of 0.5 μm and lengths stretching from 120 to 200 μm (aspect ratio ranging from 1:240 to 1:400) in posterior and anterior region respectively. Rods were tightly packed and oriented at 19° (posterior region) relative to periostracum layer and 22° (anterior region) (Fig. 1D). An earlier report demonstrated 45° of orientation for the rods in *M. edulis* in the anterior region ^25^. However, the degree of orientation of the rods in the posterior region of the shell was not in the scope of earlier studies. We further investigated the morphology of the rods in more detail using high-magnification SEM imaging (Fig. 3). We noticed the presence of nanometer-sized pores with a diameters of about 40-50 nm. These were randomly distributed throughout the length of the rods. Investigating cracked rods, we noticed that in fact these pores are opening of channels extending at around 1μm in length within the core of the calcified rods, mostly present at the anterior regions close to the ventral side.

**Fig. 3.**
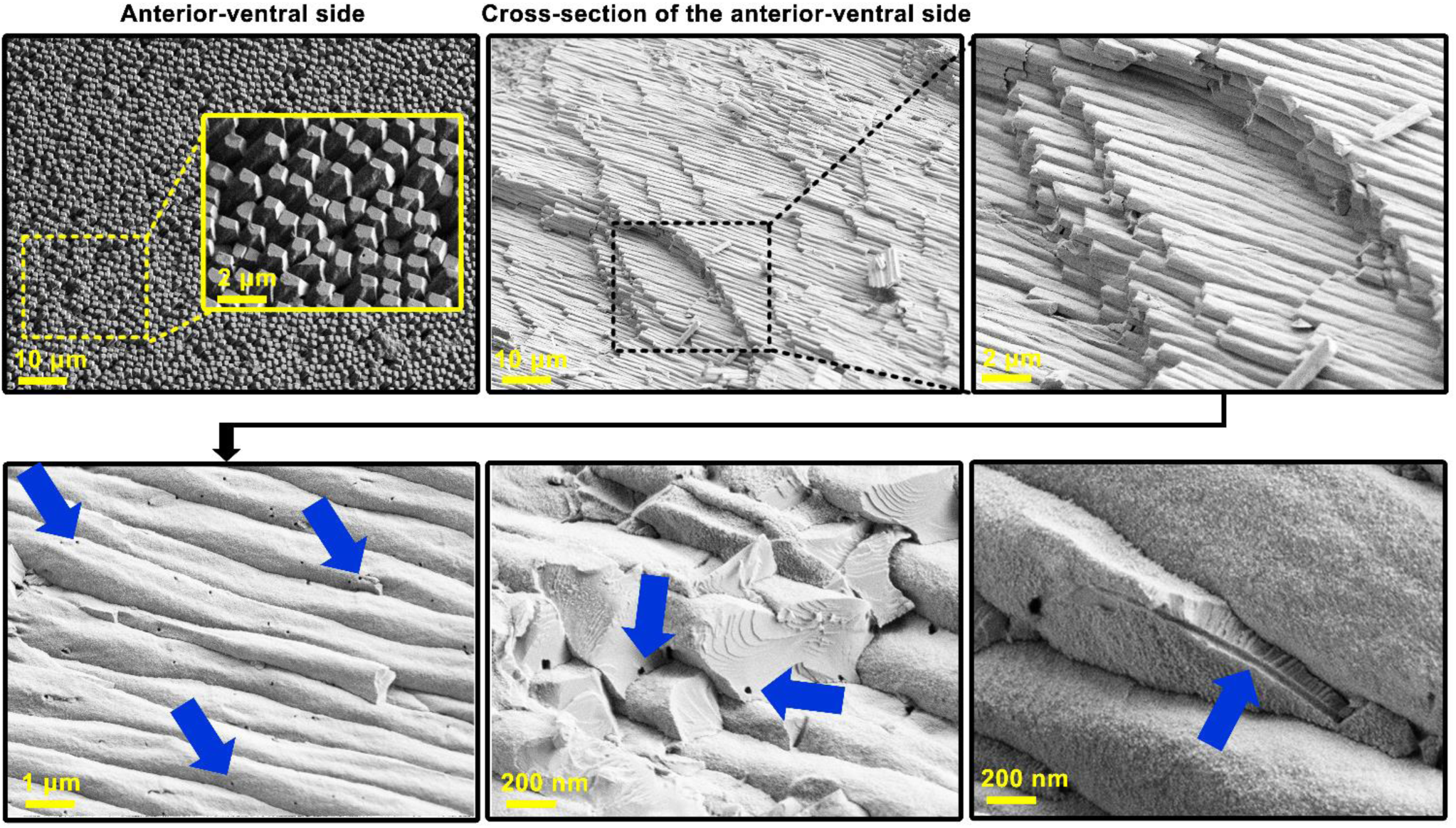
Characterization of the calcified rods in the oblique prismatic layer located at the anterior-ventral side of the shell. High magnification SEM images revealed the presence of pores with diameters of 40-50 nm forming openings of channels that extend through the rods. Blue arrows indicate the opening of the pores. Fracturing the rods also shows also that the channels extended through the rods.

### 3.5. Mechanical properties

To gain additional insight into the mechanical properties of the shell, we performed nanoindentation mapping on longitudinal and transversal cross-sectional cuts of the two identical shell originating from the same mussel (Fig. 4A, B, and fig. S2A). Probing the longitudinal cut exhibited a gradient of mechanical properties. Moving from anterior to the posterior region we collected the profile at nine positions, each three millimeters apart from each other and noticed an increase in the stiffness value of about 53 ±5.9 GPa to 80 ±16.5 GPa. For the same set of experiment, the calculated hardness showed values increasing from 1.5 GPa at the anterior region to around 4 GPa at the posterior region. We then performed nanoindentation on the other shell, in which we made a transversal cut corresponding to position five of the longitudinal cut and performed nanoindentation at five different positions (Fig. 4C, D, and fig. S2B). Moving laterally, Young’s modulus was ranging from 60 to 66 GPa to and hardness value ranging from 2.5 to 3 GPa.

**Fig. 4.**
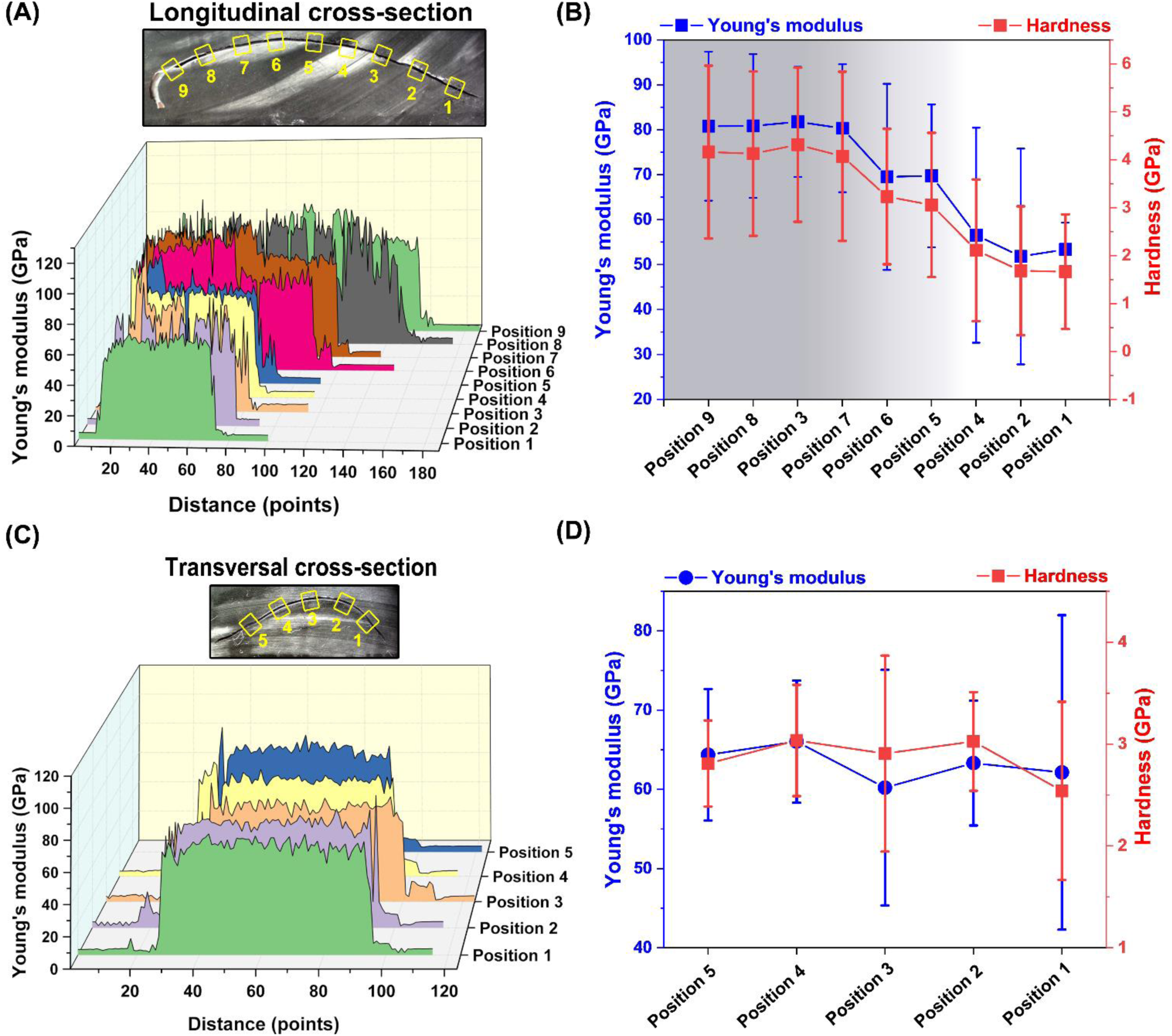
Mechanical properties derived from nanoindentation experiments performed on the longitudinal and transversal cross-section of the shell. (A) Profile of Young’s moduli obtained at nine different positions across longitudinal cross-section and (B) average Young’s modulus and hardness values for each position. (C) Profile of Young’s moduli obtained at nine different positions across transversal cross-section and (D) average Young’s modulus and hardness value for each position.

### 3.6. Organic-rich interface

By studying fracture surfaces of the hydrated oblique prismatic layer, we noticed the presence of an organic adhesive matrix filling the interfaces between the rods (Fig. 4A). Most importantly, we noticed that the gel-like adhesive matrix not only oozed out from the interfaces of the rods but also from the openings of the nano-sized pores. Similar gel-like organic matrix have been observed in fractured shell of *Atrina rigida* and *Pinctada margaritifera* ^4^.This provided the basis to hypothesize that the observed nano-sized pores may have a function in transporting matrix material during the biomineralization and adhesion of the rods. In addition, we identified regions in which adhesive organic matrix formed filaments, bridging cracks while fracturing. It is noticeable that filaments are distinctly oriented perpendicular to the direction of the crack (Fig. 4A), suggesting resistance to fracture by dissipating energy through the formation of nano-sized filaments which deflects the cracks into regions in which propagation becomes more difficult.

In order to investigate the nature of the matrix and the compositions in more detail and compare to earlier studies, we performed FTIR experiments in transmission mode. Fig. 5B and C illustrate FTIR spectra of the bulk shell powder, containing soluble and insoluble organic remains after decalcification (either using 1M EDTA or 5% v/v acetic acid). Spectra strongly exhibit the presence of organic-rich-matrix and calcium carbonate (Table S1). The organic-rich-matrix exhibits three main FTIR band ranges ^27–29^. At 834-899 cm^-1^ stretching vibrations of the C-O-C and C-C bonds corresponded to α-chitin as expected. In the range of 1100-1700 cm^-1^ multiple bands showed up, mainly corresponding to amide (I, II and III) bands of protein (C=O, C-N, N-H, C-N and C-C vibrations) and α-chitin (CH_2_, OH, and C=O vibration). Finally, the 2962 cm^-1^ band corresponds to amide B band of α-chitin.

**Fig. 5.**
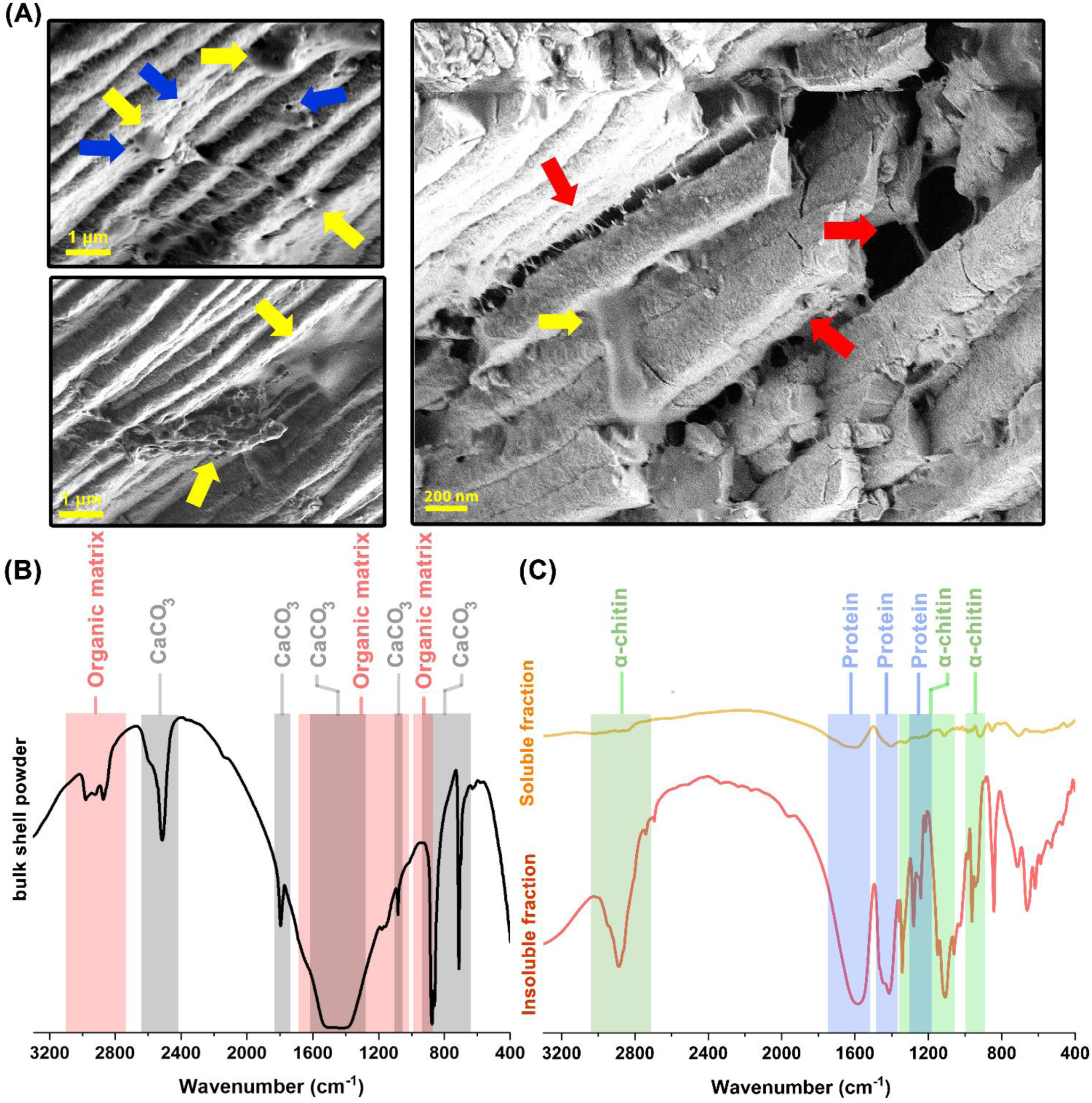
(A) Environmental SEM images from the fracture surface of the hydrated oblique prismatic layer, illustrating organic adhesive matrix filling the interfaces between the calcified rods also regions in which the organic-rich matrix bridges the crack by forming filaments. Yellow arrows show a gel-like adhesive matrix that oozes out from the interfaces of the rods but also from the openings of nano-sized channels shown by blue arrows. Red arrows show the filaments bridging the cracks and they are oriented perpendicular in the direction of crack propagation. (B) FTIR spectra of the bulk shell powder. (C) FTIR spectra of the extracted organic matrix after decalcification.

### 3.7. Protein in the matrix

It has become more and more clear that proteins play crucial roles during mineralization and assembly of marine shells. Proteins (small quantities of less than 1 %) along with the chitin network have been shown to provide structural integrity, functioning as soft, energy-dissipating matrix by hindering crack propagation through the interfaces and increasing the toughness (Feng et al., 2017; Zhang et al., 2012; Bram et al., 2012; Fabio and M., 2012). Therefore, we set to explore proteins extracted from the shell. Fig. 6A illustrates sodium dodecyl sulfate-polyacrylamide gel electrophoresis (SDS-PAGE) of the extracted proteins. Proteins were extracted after decalcification either using 1M EDTA or 5% v/v acetic acid. Presence of more protein extracts was noticeable in the insoluble fractions than soluble fractions obtained independent of extraction method. Clear bands, one at 100 kD and the other one at 45 kD in the insoluble extracts were noticeable. However, we did not observe similar bands in the soluble fraction. Further, 100 kD and 45 kD were subjected to in-gel trypsin digest (Fig. 6B) for de novo sequencing. Ten fragments from tryptic digest of 100kD protein and six from 45kD were selected, which had the strongest peak as well as the highest quality factor. Fig. 6C and D illustrates product ion spectrum and identified peptides with de novo sequencing for the corresponding isolated tryptic fragments. To identify possible hits to existing protein sequences, resulted monoisotopic masses cross-referenced using MASCOT search engine (Matrix Science, London, UK; version 2.1) against NCBI server (http://www.ncbi.nlm.nih.gov). In addition to that, identified de novo sequences cross-checked manually against BLASTp, Swiss-Prot and UniProt. However, we did not find any significant hits. Even though it would be too early to draw any conclusion about the nature of the intact proteins using these identified short fragments, but analyzing the amino acid composition Fig. S3 showed 100 kD protein might be rich in glycine, serine, asparagine, alanine, proline and arginine. However, 45 kD proteins mostly contained alanine, glycine, glutamine, lysine, serine, tryptophan and tyrosine.

**Fig. 6.**
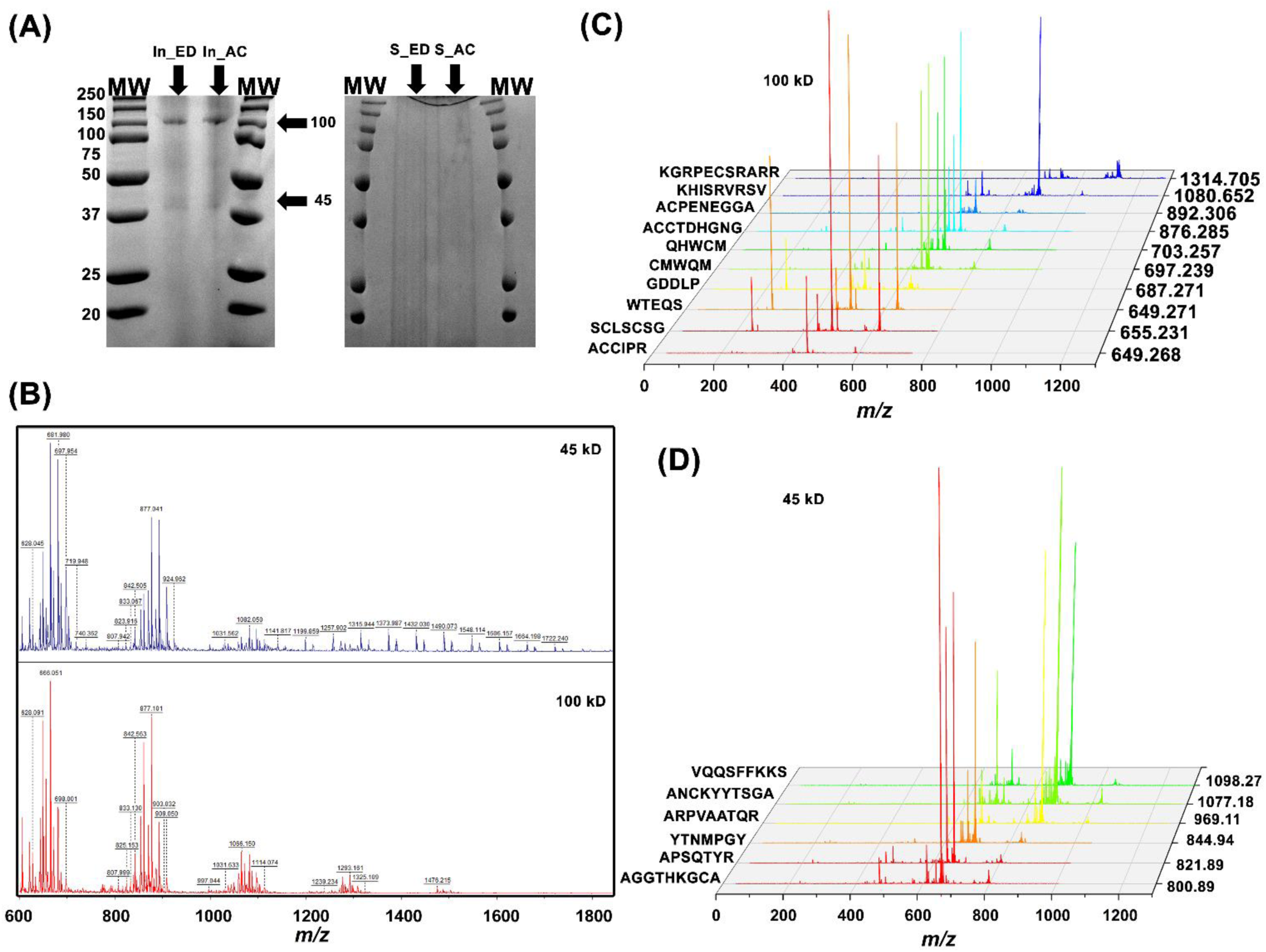
(A) SDS-PAGE of the soluble and insoluble protein extracted after decalcification of the shell using 1m EDTA pH 8 (marked as S_ED and In_ED) and 5% v/v acetic acid (marked S_AC and In_AC). Molecular weight markers are indicated as MW. (B) Mass spectrum of fragments produced from tryptic digestion of 100 kD (In_ED) and 45 kD (In_ED) band. (C) MS/MS spectrum of ten peptides obtained from 100kD (IN-ED) digest with m/z ranging from 650-1315 and their corresponding sequences. (D) MS/MS spectrum of six peptides from 45 kD (IN-ED) digest with m/z ranging from 800-1100 and their corresponding sequences.

## 4. Conclusion

To our knowledge, this is the first detailed study on the structure of *M. edulis* L. shell, commonly known as Finnish blue mussel found in the archipelago of SW-Finland (Northern Baltic Sea). We employed various techniques to investigate in detail, both inorganic mineralized components as well as the organic-rich matrix of *M. edulis* L. Being closely related to *Mytilus edulis* we found a very similar ultra-architecture across the shell. However, we also found differences, for instance in the thickness and degree of orientation of the layers. Using high resolution SEM imaging and probing the calcified rods in the oblique prismatic layer, we found pores with a diameters of 40-50 nm and lengths up to a micrometer extending through the rods. These channels were only found in the anterior ventral part which lacks the nacreous part. We also noted that the matrix material was highly fluid, oozing out from the cracked surfaces. The fluidity of the matrix and the presence of the channels in the more newly formed parts of the shell, leads us to suggest that the channels might have a role for transport of the fluid organic matrix. Our results also show that the shell exhibits a gradient of mechanical properties. Moving from anterior to posterior region, we noticed an increase of both stiffness and hardness. This likely attributes to the fact that each region has quite distinct morphology as they are in the different developmental stage. At the posterior region, the biomineralized layer grows and developed earlier, hence this resulted in thicker and more compact motifs with noticeable differences in the degree of orientation than the anterior region. We hypothesize that this could possibly be a generic characteristic for other closely related taxa of mussels including *M. edulis* as this has not been in the scope of earlier studies to our knowledge. This finding could provide inspiration for designing lightweight load-bearing materials in which a gradient of mechanical properties is crucial. There is growing evidence suggesting that the organic-rich interface largely governs both intrinsic and extrinsic toughening in such materials. In addition to that, their substantial importance for controlling both nucleation and inhibition of the crystal growth and the adhesion of the rod cannot be ignored. We identified chitin and protein as the main component of the organic-rich interfacial matrix. We found protein-chitin-rich interface forms less than 1% volume fraction of the shell. However sufficient to work in synergy with their stiff surrounding architecture to provide non-linear deformation upon initiation of the cracks. This results in the propagation of the cracks into conformations that requires considerable energy before undergoing catastrophic failure with elongated filaments of chitin-protein, bridging crack at the interface of the calcified rods. Proteomic analysis showed two abundant proteins with sizes around 100 kD and 45 kD. In future, a combination of high-throughput RNA-sequencing, advanced proteomics analysis, and X-ray crystallography could be applied to identify the full-length sequences of these proteins and find possible sequence or structural homology to other known proteins. Eventually, this could be combined with molecular dynamic simulations to accelerate our understanding of their key molecular interactions with chitin and CaCO_3_ and their colloidal complexation toward structure formation.

## Acknowledgments

We acknowledge Aalto University Nano-microscopy Center (Espoo) for their generous microscopy time. The work was performed within the Academy of Finland Center of Excellence Programme (2014–2019) and Academy of Finland projects 264493, 259034, and 317395. Chenghao Li is acknowledged for help during the synchrotron measurements at BESSY, Fabian Zemke for assistance with x-ray scattering data evaluation and Petra Leibner for performing the nanoindentation experiments.

## Author Contributions

P.M and W.W designed and conducted the experiments. M.B.L supervised the project. All Authors contributed in writing the manuscript.

## Competing financial interests

The authors declare no competing financial interest.

