## Supplementery information for "Analysis of Finnish blue mussel (*Mytilus edulis* L.) shell: Biomineral ultrastructure, organic-rich interfacial matrix and mechanical behavior"


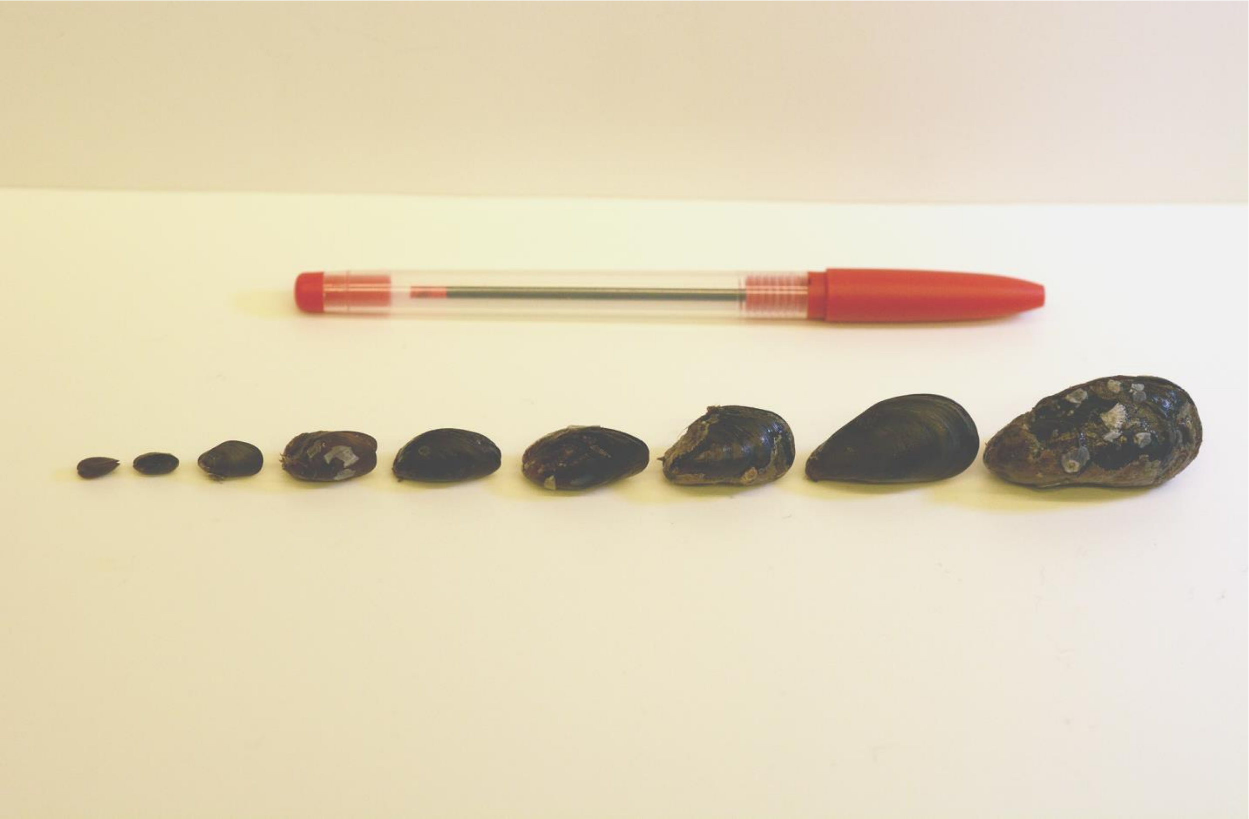

Supplementary figure 1. Illustrating *Mytilus edulis* L. collected from the archipelago of SW-Finland, northern Baltic Sea at a different stage of maturity.

| Table S1. FTIR pick assignment | | |
|---|---|---|
| **Protein** | | |
| **Wavenumber ($cm^{-1}$)** | **Bending or/or and stretching vibration** | **Ref.** |
| 1200-1700 | Amide I Band: stretching vibration of the C=O bonds in the random coil or/and helix in protein (1600 $cm^{-1}$)<br>Amide I Band: stretching vibration of the C-N bonds<br>Amide II Band: in-plane bending vibration of N-H bonds in the random coil or/and helix in protein (1484 $cm^{-1}$)<br>Amide II Band: stretching vibration of C-N bonds<br>Amide II Band: stretching vibration of C-C bonds<br>Amide III Band: resulted mainly from a mixture of C–N vibration (1260 $cm^{-1}$ )<br>Amide III Band: resulted from some N–H vibration (1232 $cm^{-1}$) | [1] |
| **Chitin** | | |
| **Wavenumber ($cm^{-1}$)** | **Bending or/or and stretching vibration** | **Ref.** |
| 800-900 | Stretching vibration of the C-O-C (834 $cm^{-1}$) bond in alpha chitin<br>Stretching vibration of the C-C (899 $cm^{-1}$) bond in chitin | [1] |
| 1100-1700 | Amide I Band stretching vibration of the C=O bonds in the random coil or/and helix in alpha-chitin (1660 $cm^{-1}$)<br>Amide II band δ: bending vibration of the CH2 in alpha-chitin (1550 $cm^{-1}$)<br>δ: bending vibration of the CH2 in alpha-chitin (1460 $cm^{-1}$)<br>δ: bending vibration of the OH hydrogen bonded to C=O in alpha-chitin (1620$cm^{-1}$)<br>Amide III bands of chitin alpha-helix (1280 $cm^{-1}$)<br>Amide III bands of chitin alpha-helix (1330 $cm^{-1}$)<br>Amide III bands of chitin alpha-helix (1380 $cm^{-1}$) | [1] |
| 2900-3000 | Amide B band of chitin alpha-helix (2962 $cm^{-1}$) | [2] |
| **$CaCo_3$** | | |
| **Wavenumber ($cm^{-1}$)** | **Bending or/or and stretching vibration** | [4] |
| 700-900 | $V_4$: symmetric bending vibration of the $CO_3$ (725 $cm^{-1}$)<br>$V_2$: asymmetric bending vibration of the $CO_3$ (874 $cm^{-1}$) | |
| 1000-1100 | $V_1$: symmetric bending vibration of the $CO_3$ (1090 $cm^{-1}$) | |
| 1300-1500 | $V_3$: asymmetric bending vibration of the $CO_3$ (1409 $cm^{-1}$) | |
| 1700-1900 | $V_1$ + : $V_4$ asymmetric bending vibration of the $CO_3$ (1805 $cm^{-1}$) | |
| 2400-2600 | Bending vibration of $HCO_2^-$ (5230 $cm^{-1}$) | [5] |

Table S1. Assignments of picks observed in the FTIR.

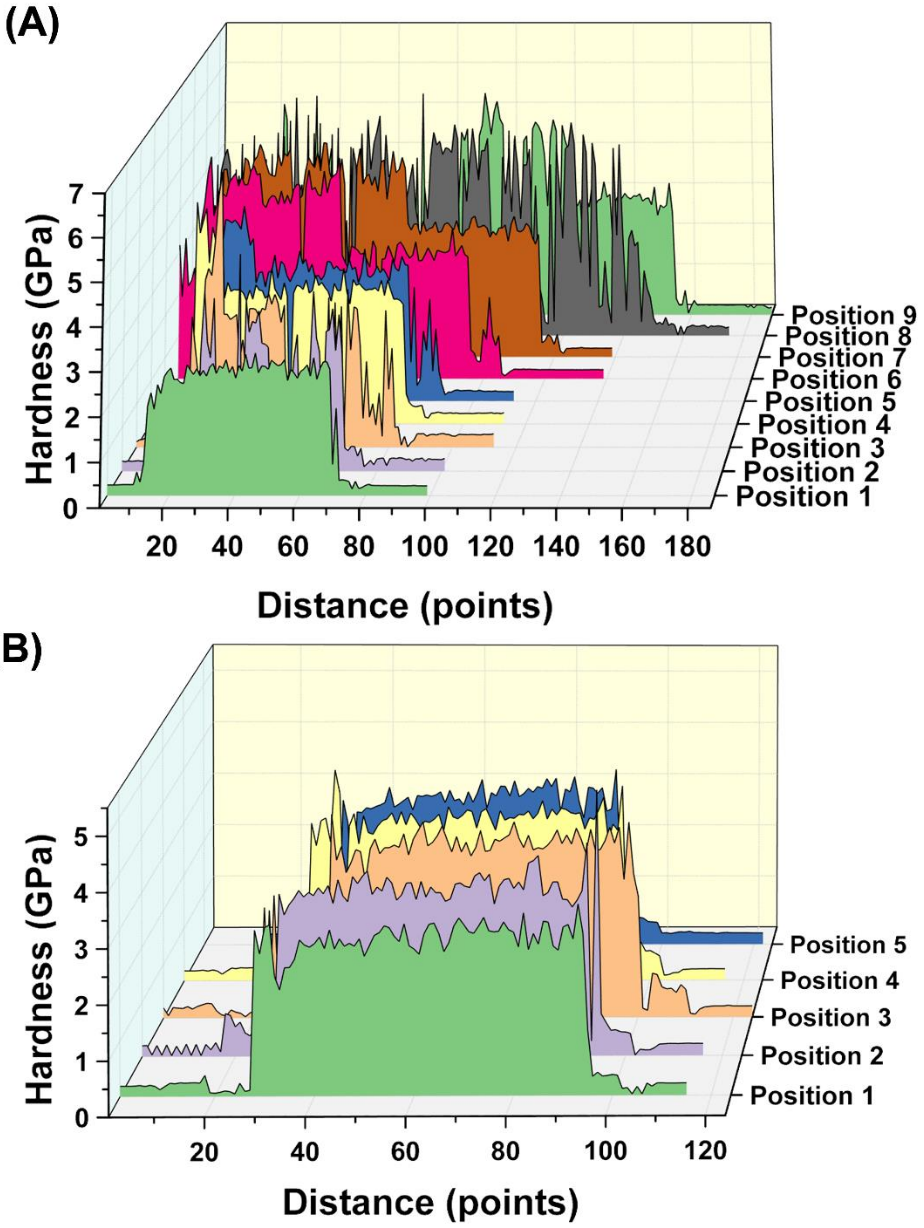


Supplementary figure 2. Extracted hardness profile derived from nanoindentation experiment performed on the Longitudinal (A) and transversal (B) cross-sectional cuts.

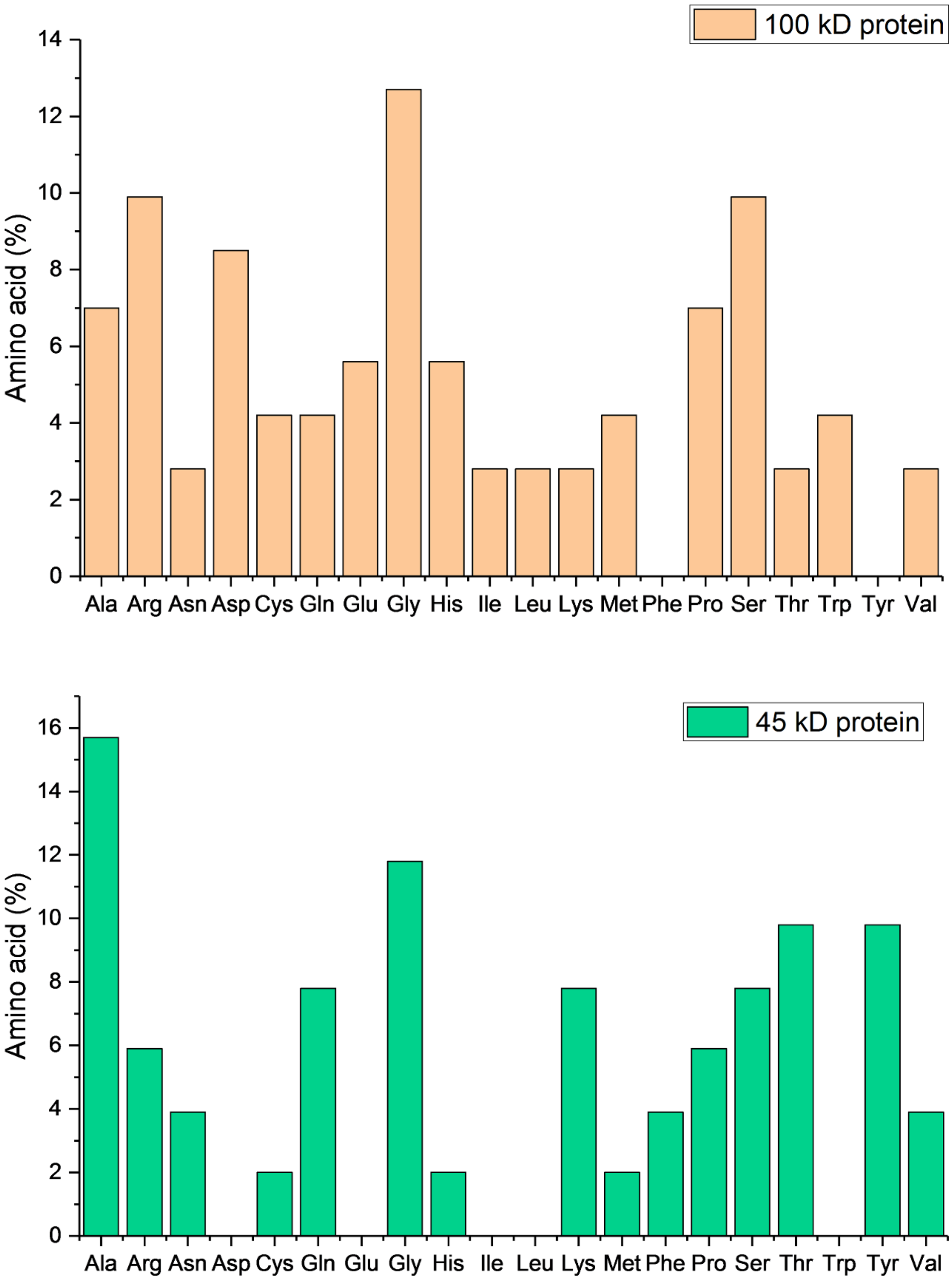


Supplementary figure 3. The amino acid composition of identified peptides with de novo sequencing for 100 kD and 45 kD protein.
